# Impact of a resistance gene against a fungal pathogen on the plant host residue microbiome: the case of the *Leptosphaeria maculans-Brassica napus* pathosystem

**DOI:** 10.1101/2020.01.13.903765

**Authors:** Lydie Kerdraon, Matthieu Barret, Marie-Hélène Balesdent, Frédéric Suffert, Valérie Laval

## Abstract

Oilseed rape residues are a crucial determinant of stem canker epidemiology, as they support the sexual reproduction of the fungal pathogen *Leptosphaeria maculans*. The aim of this study was to characterise the impact of a resistance gene against *L. maculans* infection on residue microbial communities and to identify micro-organisms interacting with this pathogen during residue degradation. We used near-isogenic lines to obtain healthy and infected host plants. The microbiome associated with the two types of plant residues was characterised by metabarcoding. A combination of linear discriminant analysis and ecological network analysis was used to compare the microbial communities and to identify micro-organisms interacting with *L. maculans*. Fungal community structure differed between the two lines at harvest, but not subsequently, suggesting that the presence/absence of the resistance gene influences the microbiome at the base of the stem whilst the plant is alive, but that this does not necessarily lead to differential colonisation of the residues by fungi. Direct interactions with other members of the community involved many fungal and bacterial ASVs (amplicon sequence variants). *L. maculans* appeared to play a minor role in networks, whereas one ASV affiliated to *Plenodomus biglobosus* (synonym *Leptosphaeria biglobosa*) from the *Leptosphaeria* species complex may be considered a keystone taxon in the networks at harvest. This approach could be used to identify and promote micro-organisms with beneficial effects against residue-borne pathogens, and more broadly, to decipher the complex interactions between multi-species pathosystems and other microbial components in crop residues.

## Introduction

Plants support a large number of micro-organisms, and the assembly and structuring of this microbial community is dependent on many factors, such as the type of plant, the organ considered (Comby et al., 2016; Grudzinska-Sterno et al., 2016) and its age (Wagner et al., 2016). Many of these micro-organisms are considered beneficial (e.g. plant growth-promoting bacteria), whereas others are pathogenic and decrease the yield and quality of agricultural produce. The pathobiome is the subset of microbial communities consisting of a pathogen and the organisms that can influence that pathogen or be influenced by it (Jakuschkin et al., 2016; Vayssier-Taussat et al., 2014). The role of the microbiota in the plant’s response to a disease or in the pathogenicity of a fungal pathogen is currently being studied in various pathosystems, but remains poorly understood (Cho and Blaser, 2012; Jakuschkin et al., 2016; Lebreton et al., 2019).

Stem canker is a widespread disease of oilseed rape (*Brassica napus*), the main causal agent of which is the ascomycete *Leptosphaeria maculans*. This fungus has a complex life cycle. It enters the leaves through stomata (Hammond and Lewis, 1987), leading to the development of leaf spots (Travadon et al., 2007). The fungus then progresses through the xylem, from the leaf spots to the base of the petiole and into the stem (Hammond and Lewis, 1987; Hammond et al., 1985). In Europe, *L. maculans* begins its necrotrophic phase in late spring, causing the “crown canker” (or “stem canker”) at the stem base (West et al., 2001), leading to step breakage yield losses. The fungus continues its life cycle as a saprophyte on the infected stem bases left on the soil at harvest, on which it reproduces sexually and survives the intercropping period, sometimes over a period of several years (Petrie and Lewis, 1985; Baird et al., 1999; West et al., 2001; Fitt et al., 2006). Control strategies are mostly based on improving plant immunity by combining quantitative, polygenic resistance sources, and major resistance genes (*Rlm* genes) matching fungal effectors acting as avirulence (*AvrLm*) determinants (Delourme et al., 2006).

Another *Leptosphaeria* species, *Leptosphaeria biglobosa*, has a similar life cycle to *L. maculans* and colonises oilseed rape plant tissues in a similar manner. This fungus is present in all countries except China (Liu et al., 2014), but its agronomic impact is considered negligible relative to that of *L. maculans. L. biglobosa* was recently renamed *Plenodomus biglobosus* (De Gruyter et al., 2013), but this nomenclature has not yet been adopted as the standard among plant pathologists working on oilseed rape stem canker (Dutreux et al., 2018). For the sake of consistency, we use ‘*P. biglobosus*’ here, because the genus name ‘*Plenodomus*’ is derived from the taxonomic assignment of metabarcoding sequences.

Crop residues are an essential ecological niche for both *L. maculans* and *P. biglobosus*, and the severity of stem canker disease is generally correlated with the number of ascospores ejected by the residues of the previous crop (McGee and Emmett, 1977; Petrie, 1995). Residues also constitute a compartment rich in micro-organisms, in which transfers occur between organisms from the plant and micro-organisms from the soil (Kerdraon et al., 2019a). These micro-organisms may interact with *L. maculans*, or be influenced by its presence, as recently established for *Fusarium* sp. in maize (Cobo-Díaz et al., 2019) and *Zymoseptoria tritici* in wheat (Kerdraon et al., 2019b). Culture-dependent approaches have shown that some fungal species (e.g. *Alternaria* spp., *Trichoderma* spp., *Chaetomium* sp., *Gliocladium* sp.) are present together with *L. maculans* on buried oilseed rape residues (Naseri et al., 2008). The characterisation of interactions between pathogens and other micro-organisms can open up new opportunities for managing residue-borne diseases (Poudel et al., 2016).

An initial description of the microbiota associated with *L. maculans* during its saprophytic/reproduction stage on oilseed rape residues was obtained by culture-independent approaches (Kerdraon et al., 2019c). *L. maculans* and *P. biglobosus* were both found to be highly represented in the metabarcoding data set. However, questions remain about the influence of these species on the fungal and bacterial communities and the effect of *L. maculans* on the composition and evolution of the residue microbiome. We addressed these questions with a pair of near-isogenic oilseed rape lines with the same genetic background, differing by the presence or absence of the *Rlm11* resistance gene: ‘Darmor’ and ‘Darmor- *Rlm11*’. The major gene *Rlm11* was introgressed from *Brassica rapa* into the French cultivar Darmor following an interspecific cross and a series of backcrosses to Darmor (Balesdent et al., 2013). Population surveys showed that more than 95% of French *L. maculans* isolates were avirulent toward *Rlm11* (Balesdent et al., 2013). In addition, Darmor is characterised by a high level of quantitative, polygenic resistance. This context constituted the perfect framework for a metabarcoding approach to investigate, under field conditions, how the presence of one efficient major resistance gene, and hence the lack or low levels of *L. maculans*, affected the plant residue microbiome, including other plant pathogens of oilseed rape.

## Results

### Overall diversity

The impact of the *Rlm11* resistance gene on the oilseed rape residue microbiome was assessed by analysing the composition of fungal and bacterial communities of 120 residue samples (60 from Darmor and 60 from Darmor-*Rlm11* residues) obtained during four different sampling periods (July, i.e. just after harvest; October, i.e. just after sowing of the subsequent wheat crop; December; and February). Metabarcoding analysis identified 610 bacterial ASVs and 335 fungal ASVs. Fungal diversity (Shannon index) and observed richness were higher in Darmor-*Rlm11* residues than in Darmor residues in July, but did not differ significantly between these two types of residues on subsequent sampling dates, over which it remained relatively constant (Suppl. Figure 2 and 3). Bacterial diversity, observed richness and Pielou’s evenness increased from July to October for both cultivars, and then slightly decreased from December to February. Fungal diversity remained constant throughout the experiment, for both cultivars.

**Figure 1.**
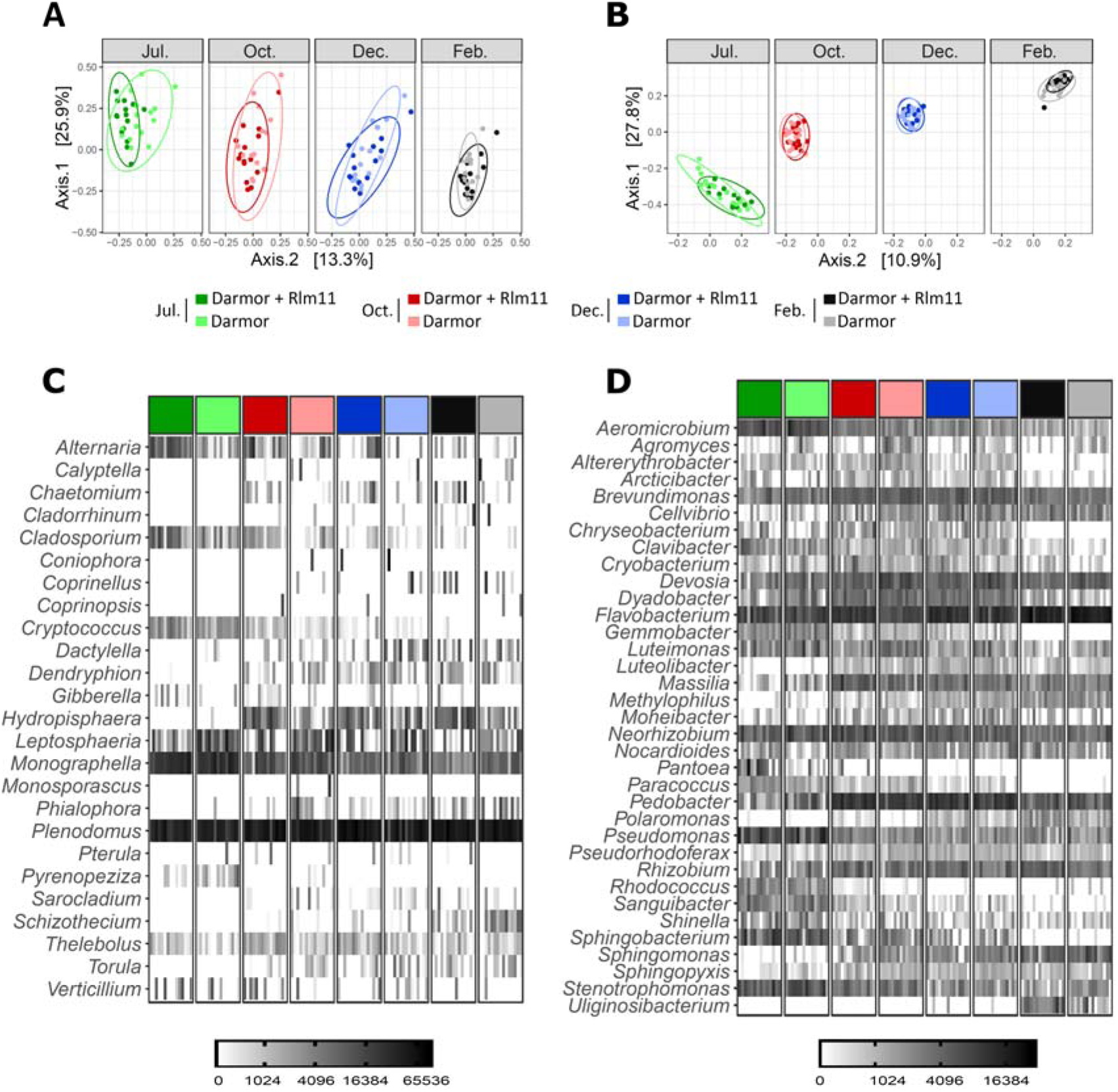
Effect of the presence of the *Rlm11* resistance gene and of sampling date (July, October, December and February) on fungal (A, C) and bacterial (B, D) communities originating from 120 samples of oilseed rape residues. (A, B) Visualisation of compositional distances between samples through multidimensional scaling (MDS) based on Bray-Curtis dissimilarity matrix. MDS analysis was performed on all samples together and was faceted according to the sampling date. Each data point corresponds to one sample of oilseed rape residues. The colours of the points distinguish between sampling dates (July: green; October: red; December: blue; February: grey) and cultivar (Darmor: light hues; Darmor-*Rlm11*: dark hues). (C, D) Diversity and predominance of the 25 most abundant (25/62 – excluding unclassified genera) fungal genera (A) and the 35 most abundant (35/134 – excluding unclassified genera) bacterial genera (B) distributed in all samples, distinguishing between the different experimental conditions. The colours used to distinguish samples are as in (A) and (B).

**Figure 2.**
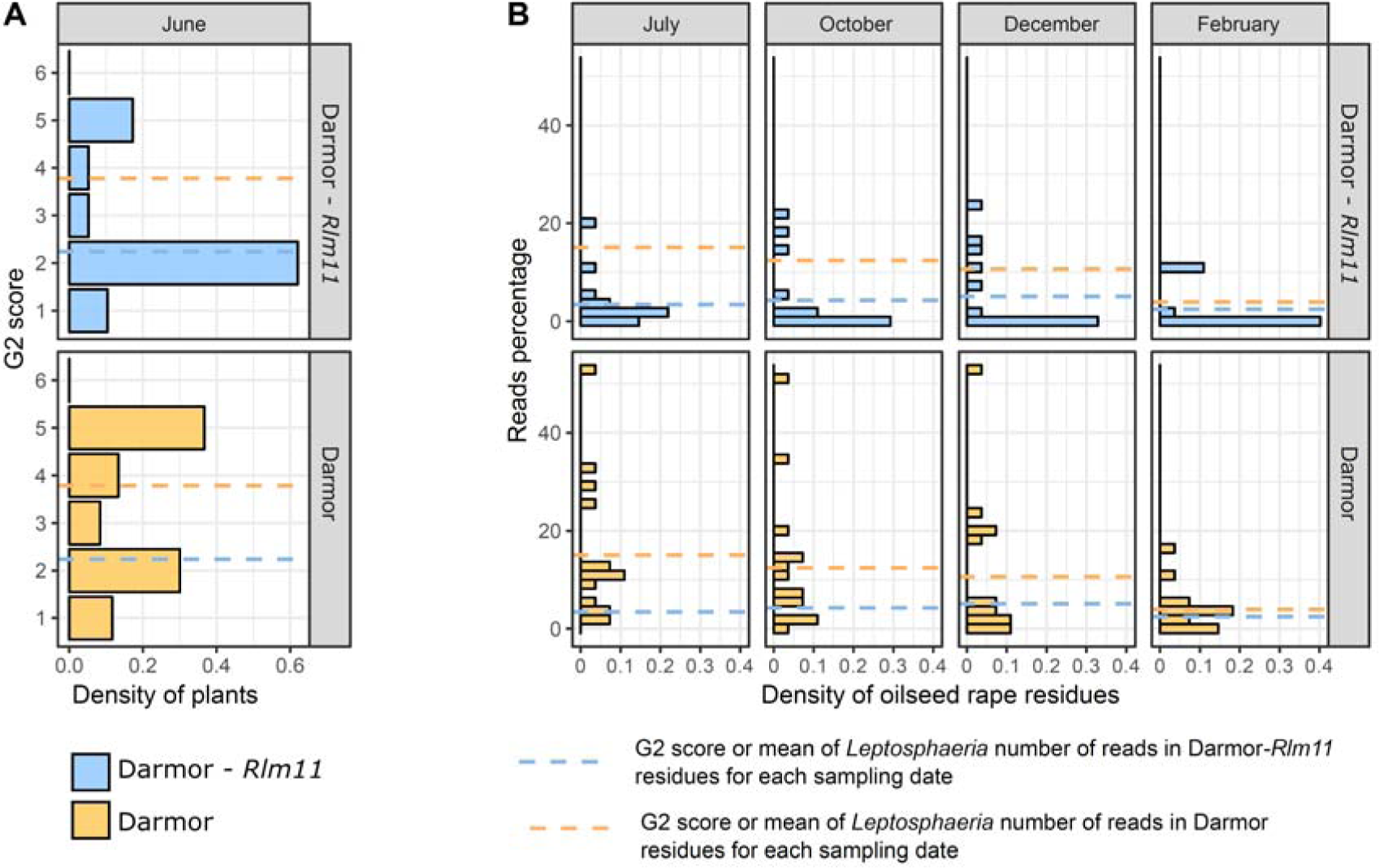
Influence of the presence of the *Rlm11* resistance gene and sampling date (July, October, December and February) on (A) the stem canker ‘G2 score’ (proxy for disease severity, estimated for 60 plants of Darmor and Darmor-*Rlm11*) and (B) the percentage of reads corresponding to *Leptosphaeria maculans*. (A) Percentage of plants with each G2 score for the two cultivars. The dashed lines correspond to the G2 score for each of the two cultivars (Darmor: yellow; Darmor-*Rlm11*: blue). (B) Percentage of reads affiliated to *L. maculans* in the 15 samples for each condition. The dashed lines correspond to the mean read percentage, for each of the two cultivars, at each sampling date (Darmor: yellow; Darmor-*Rlm11*: blue).

**Figure 3.**
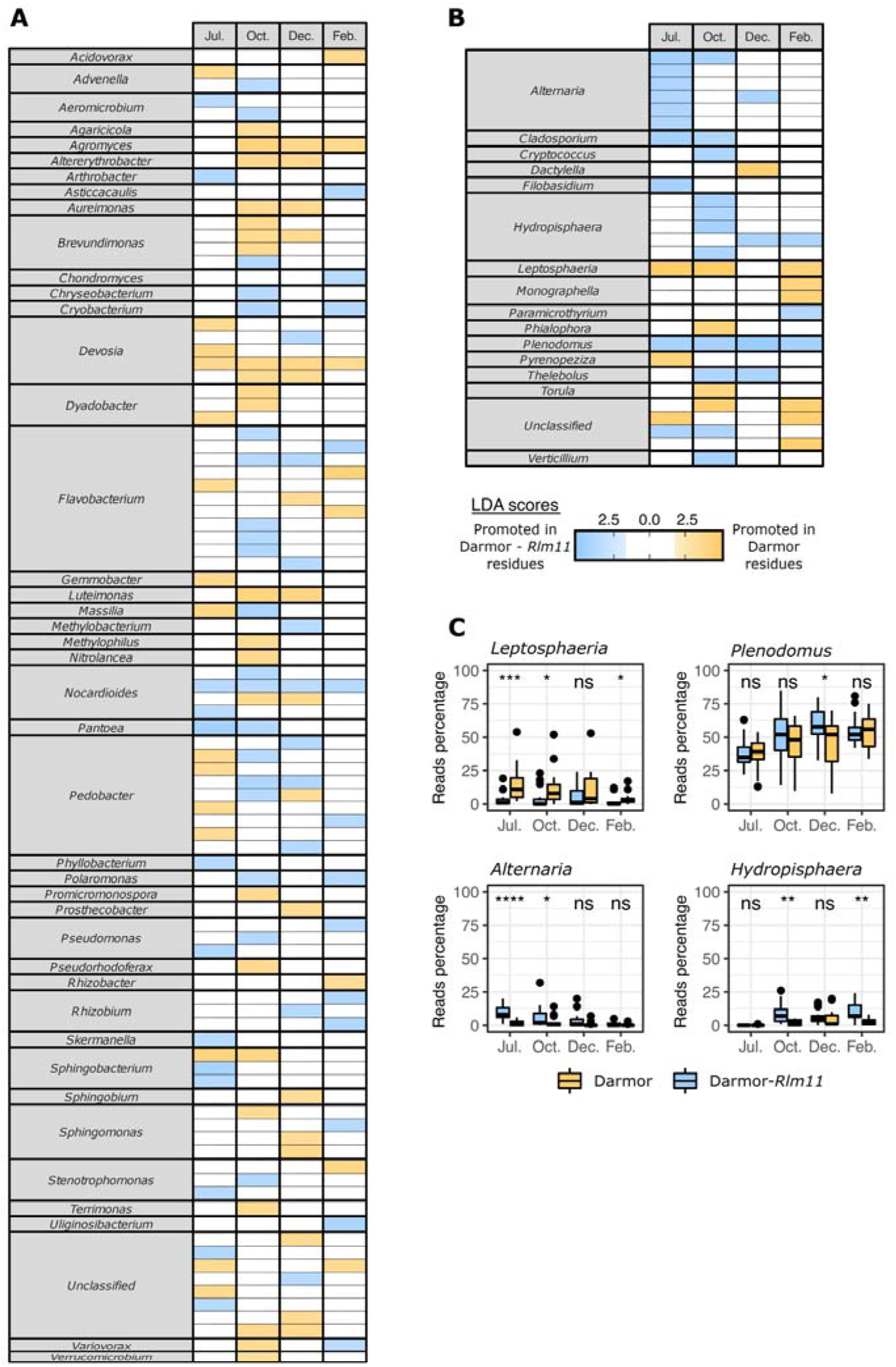
Impact of the presence of the *Rlm11* resistance gene on microbial communities in oilseed rape residues. (A, B) Significant differences in the predominance of bacterial (A) and fungal (B) ASVs between the Darmor samples (orange) and the Darmor-*Rlm11* samples (blue) obtained in linear discriminant analyses (LDA). Only ASVs with *p*-values < 0.05 for the Kruskal-Wallis test and LDA scores > 2 were retained for the plot. (C) Relative abundances of *Leptosphaeria, Plenodomus, Alternaria* and *Hydropisphaera* by host cultivar (Darmor and Darmor-*Rlm11*) and sampling date (July, October, December and February). All ASVs were grouped by genus (*Alternaria*: 8 ASVs; *Hydropisphaera*: 8 ASVs; *Leptosphaeria*: 13 ASVs; *Plenodomus*: 81 ASVs).

### Microbial community structure

The Bray-Curtis index (beta diversity analysis) was used to characterise divergence in community structure according to the presence of *Rlm11* and sampling date. The presence of *Rlm11* significantly modified the structure of the bacterial (PERMANOVA, R^2^ = 0.022, *p*- value = 0.001) and fungal (PERMANOVA, R^2^ = 0.042, *p*-value = 0.001) communities. For fungi, the difference between Darmor and Darmor-*Rlm11* was significant only in July (Table 1, Figure 1). Beta diversity analysis highlighted changes in the community over time, for both fungi (PERMANOVA R^2^ = 0.178, *p*-value = 0.001) and bacteria (PERMANOVA, R^2^ = 0.37, *p*-value = 0.001). The strong divergence observed between July and the other sampling dates for both bacterial and fungal communities (Table 1) is consistent with the rapid degradation of residues after the summer (loss of almost 50% in terms of weight, Suppl. Figure 4). Some genera, such as *Cryptococcus, Alternaria, Sphingobacterium* and *Rhodococcus*, decreased in abundance or disappeared over time, whereas others, such as *Torula, Schizotecium, Sphingomonas* and *Sphingopyxis* emerged over time (**Figure 1**).

**Table 1.** Divergences of fungal (white corner) and bacterial (grey corner) assembly structures between the oilseed rape residues from Darmor-*Rlm11* and Darmor, assessed by pairwise adonis analysis. The significance of divergences was assessed by calculating a *q*-value (**: *q*< 0.01).

|  |  | Darmor-*Rlm11* |  |  |  | Darmor |  |  |  |
| --- | --- | --- | --- | --- | --- | --- | --- | --- | --- |
|  |  | Jul. | Oct. | Dec. | Feb. | Jul. | Oct. | Dec. | Feb. |
| Darmor-*Rlm11* | Jul. | - | 0.335** | 0.385** | 0.459** | 0.078 | 0.345** | 0.364** | 0.457** |
|  | Oct. | 0.201** | - | 0.104** | 0.329** | 0.286** | 0.105** | 0.132** | 0.333** |
|  | Dec. | 0.277** | 0.041 | - | 0.212** | 0.346** | 0.137** | 0.056 | 0.217** |
|  | Feb. | 0.376** | 0.096** | 0.044 | - | 0.446** | 0.339** | 0.217** | 0.069** |
| Darmor | Jul. | 0.142** | 0.202** | 0.243** | 0.337** | - | 0.276** | 0.316** | 0.435** |
|  | Oct. | 0.223** | 0.087 | 0.084 | 0.155** | 0.124 | - | 0.083 | 0.324** |
|  | Dec. | 0.254** | 0.083 | 0.048 | 0.089 | 0.171** | 0.041 | - | 0.195** |
|  | Feb. | 0.352** | 0.113** | 0.064 | 0.064 | 0.283** | 0.095 | 0.057 | - |

**Figure 4.**
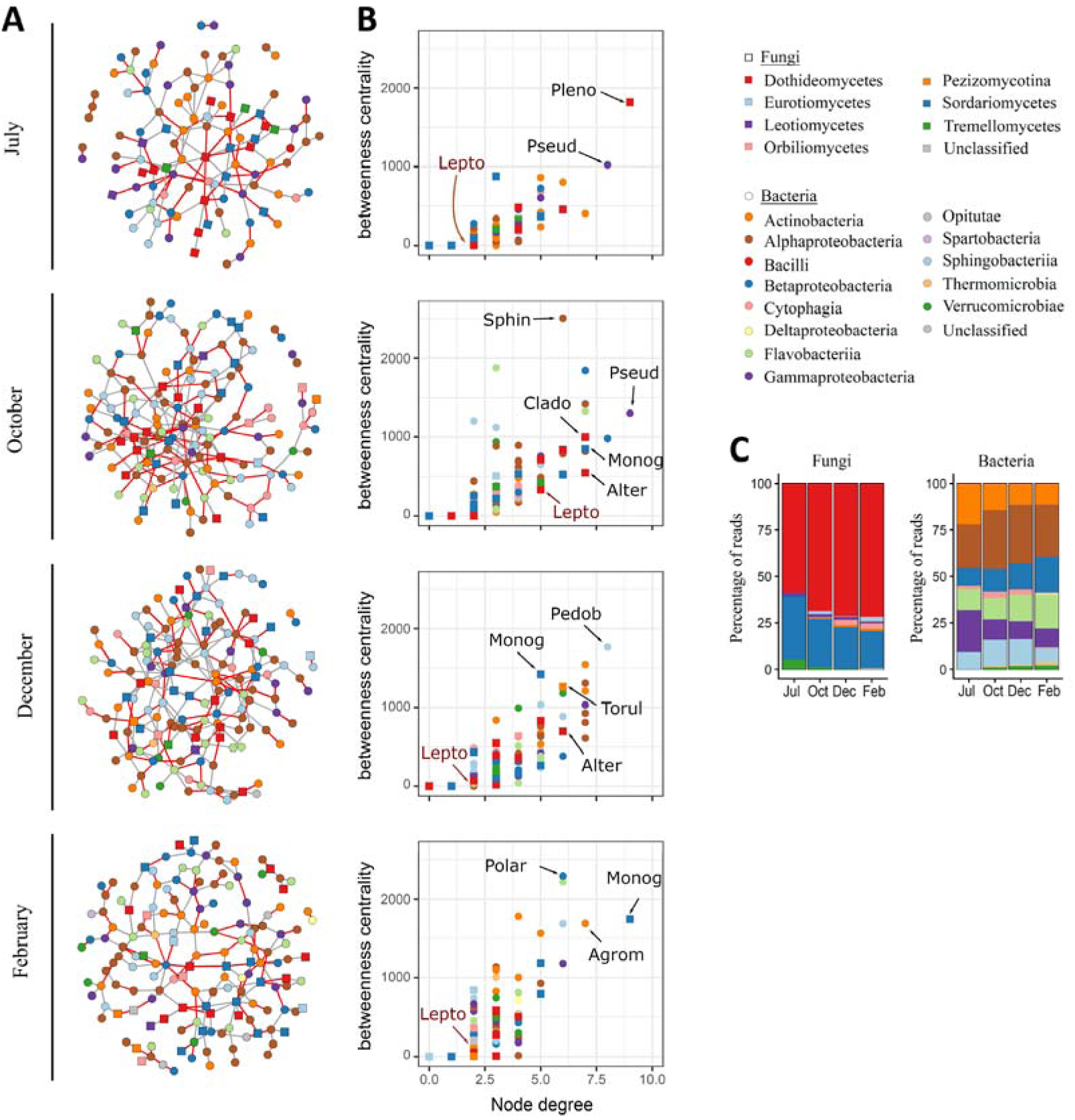
Temporal dynamics of co-occurrence networks. (A) Networks based on bacterial and fungal ASVs combined. In all networks, circles and squares correspond to bacterial and fungal ASVs, respectively, with colours representing the class. Isolated nodes are not indicated. Edges represent positive (grey) or negative (red) interactions. (B) Betweenness centrality and degree of each ASV in the networks. The place of *Leptosphaeria maculans* in the networks is indicated. Colour and shape are as in (A). The genera of the fungal and bacterial ASVs with the highest degree and centrality have been added: *Agrom(yces), Alter(naria), Clado(sporium), Lepto(sphaeria), Monog(raphella), Pedob(acter), Pleno(domus), Polar(omonas), Pseud(omonas), Sphin(gopyxis), Torul(a)*. (C) Percentage of reads associated with fungal and bacterial classes for each network. Isolated nodes are indicated. Colours are as in (A).

### Disease assessment and presence of *L. maculans* and *P. biglobosus* on residues

The G2 score, characterising stem canker severity from 0 (no disease) to 9 (all plants lodged), was obtained for the 60 oilseed rape residue samples collected in June highlighted the difference in susceptibility to *L. maculans* between Darmor (G2 = 3.78) and Darmor-*Rlm11* (G2 = 2.24). Some Darmor-*Rlm11* plants had stem canker symptoms, but the presence of *Rlm11* reduced stem necrosis due to *L. maculans* (Figure 2). Consistently, *Leptosphaeria* (probably *L. maculans*) was detected less frequently and less strongly by metabarcoding in Darmor-*Rlm11* residue samples than in Darmor residue samples. Moreover, the relative abundance (RA) of reads associated with *Leptosphaeria* decreased during residue degradation for Darmor (from 15.07 ± 14.39 in July to 3.95 ± 4.63 in February) but was lower and remained constant for Darmor-*Rlm11* (3.85 ± 6.65). The RA of ASVs affiliated to *Plenodomus* (probably *P. biglobosus*) increased over time for Darmor (from 37.10 ± 11.67 to 53.18 ± 12.62) and Darmor-*Rlm11* (from 38.47 ± 11.80 to 55.32 ± 11.66) (**Figure 3C**). The good relationship between the amount of *P. biglobosus* and *L. maculans* DNA quantified by qPCR in 33 residue samples and the corresponding sequence read numbers, as established by Spearman’s rank correlation test (ρ = 0.93; *p*-value < 0.0001 for *L. maculans*; ρ = 0.60; *p*- value < 0.001 for *P. biglobosus*; Suppl. Figure 5), justified the quantitative interpretation of metabarcoding data.

**Figure 5.**
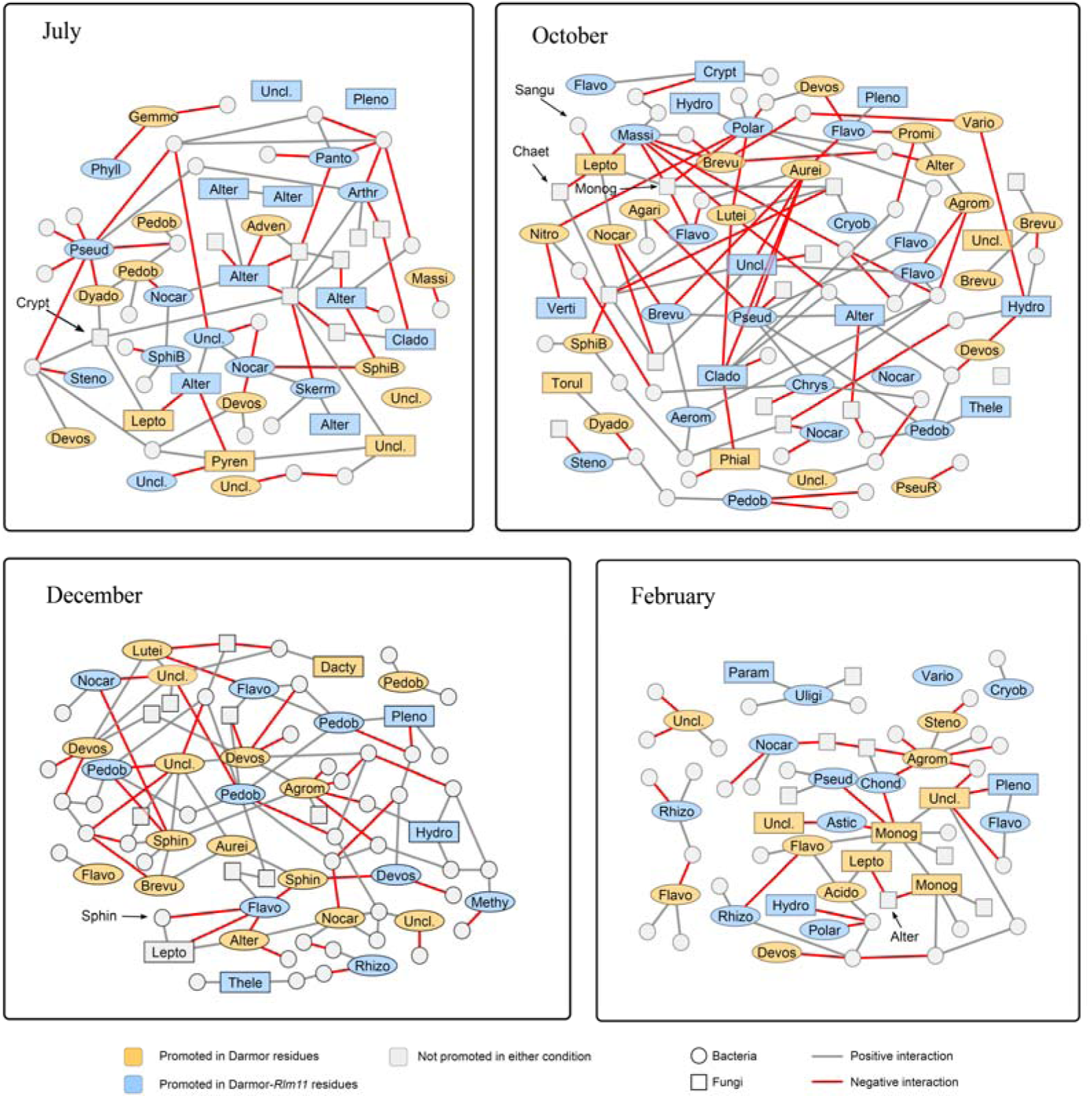
Subnetworks combining linear discriminant analyses (LDA; see Figure 3) and ecological interaction networks (see **Figure 4**), focusing on bacterial and fungal ASVs identified as differential in LDA and their first adjacent nodes. Node colour indicates the results of LefSe differential analysis between Darmor (yellow) and Darmor-*Rlm11* (blue) treatments. Only genera with *p*-values < 0.05 for the Kruskal-Wallis test and LDA scores > 2 were retained for the plot. Edges represent positive (grey) or negative (red) interactions. Differential ASVs were plotted with genus name abbreviations: *Acido(vorax), Adven(ella), Aerom(icrobium), Agari(cicola), Agrom(yces), Alter(naria), Alter(erythrobacter), Arthr(obacter), Astic(cacaulis), Aurei(monas), Brevu(ndimonas), Chond(romyces), Chrys(eobacterium), Clado(sporium), Cryob(acterium), Crypt(ococcus), Dacty(lella), Devos(ia), Dyado(bacter), Flavo(bacterium), Gemmo(bacter), Hydro(pisphaera), Lepto(sphaeria), Lutei(monas), Massi(lia), Methy(lobacterium), Monog(raphella), Nitro(lancea), Nocar(dioides), Panto(ea), Param(icrothyrium), Pedob(acter), Phial(ophora), Phyll(obacterium), Pleno(domus), Polar(omonas), Promi(cromonospora), Pseud(omonas), PseuR(=pseudorhodoferax), Pyren(opeziza), Rhizo(bium), Skerm(anella), SphiB(=Sphingobacterium), Sphin(gomonas), Steno(trophomonas), Thele(bolus), Torul(a), Uligi(nosibacterium), Uncl*.*(assified), Vario(vorax), Verti(cillium)*.

### Impact of the presence of *Rlm11* on microbial community composition

Linear discriminant analysis (LDA) was used to characterise the ASVs affected by the presence of *Rlm11*. The RA of 29 fungal ASVs (16 genera, including unclassified) was affected by the presence/absence of *Rlm11* (Figure 3). In July, fungal communities differed significantly between Darmor and Darmor-*Rlm11* (Table 1), due to a higher RA of ASVs affiliated to *Alternaria, Cladosporium, Filobasidium* and *Plenodomus* in Darmor-*Rlm11* residues, and a higher RA of ASVs affiliated to *Leptosphaeria* and *Pyrenopeziza* in Darmor residues. The communities were not significantly different overall in October, December and February (Table 1), but some ASVs displayed significantly different abundances between Darmor and Darmor-*Rlm11*. For example, one ASV (the second most abundant, with 4.5% of reads) affiliated to *Plenodomus* was favoured by the degradation of Darmor-*Rlm11* residues. The genus *Hydropisphaera*, which colonised residues after July (Figure 1), was more abundant in Darmor-*Rlm11* than in Darmor residues (Figure 3). Surprisingly, the genus *Pyrenopeziza*, including *Pyrenopeziza brassicae*, the causal agent of light leaf spot disease in oilseed rape, which reproduces on oilseed rape residues (Gilles et al., 2001), was more abundant in Darmor than in Darmor-*Rlm11* in July.

The RA of 93 bacterial ASVs (43 genera, including unclassified) was affected by the presence/absence of *Rlm11* (Figure 3). Bacterial community structure differed significantly between the two types of residues in October and February (**Table 1**). The bacteria colonising residues were affected by the presence of *Rlm11*. For example, ASVs affiliated to *Sphingomonas* preferentially colonised Darmor residues in October, and *Uliginosibacterium* preferentially colonised Darmor-*Rlm11* residues in February. No significant difference in microbial communities was found between the two types of residues in July, but 28 bacteria had different abundances in Darmor and Darmor-*Rlm11*, including *Pantoea*, which was more abundant in Darmor-*Rlm11* residue samples in July.

### Direct impact of *L. maculans* on micro-organisms

#### Dynamics of ecological interaction networks associated with oilseed rape residues

Ecological interaction networks (ENA) combining bacterial and fungal datasets were built to characterise the interactions between *Leptosphaeria* (probably *L. maculans*) and the other members of the bacterial and fungal communities, at each of the four sampling dates (Figure 4). The mean number of nodes in networks (138.8 ± 22.1) increased during residue degradation. On average, 11.8 ± 5.5 nodes were isolated. Mean node degree was 2.49 ± 0.34. *Leptosphaeria* appeared to play a weak role in networks, with only a few degrees (2 to 5 only), and a low ranking for betweenness centrality (Figure 4). By contrast, one ASV affiliated to *Plenodomus* (probably *P. biglobosus*, the other oilseed rape pathogen associated with *L. maculans*), may be considered a “keystone taxa” in the July network following the definition proposed by Banerjee et al. (2018).

#### Subnetworks highlighting direct interactions between *L. maculans* and other species

The combination of ENA and LDA highlighted interactions between the ASVs affected by the presence/absence of *Rlm11*. These ASVs were mostly connected. *Leptosphaeria* interacted with as many ASVs promoted or inhibited by the presence of *Rlm11* as with ASVs not affected by the presence of this gene (Figure 5). ASVs interacting with *Leptosphaeria* but not affected by the presence of *Rlm11* had RA values that changed with the presence of *Leptosphaeria*, regardless of the cultivar. During residue degradation, *Leptosphaeria* interacted positively with ASVs affiliated to *Cryptococcus* (July), *Nocardioides* (October), *Monographella* (in October and February), *Altererythrobacter, Sphingomonas* (December) and *Acidovorax* (February), and interacted negatively with *Alternaria* (July and February), *Massilia, Sanguibacter* and *Chaetomium* (October).

## Discussion

The use of two near-isogenic lines appeared to be a relevant strategy to characterise the effect of *L. maculans* on the microbiome of oilseed rape residues. Indeed, oilseed rape residues with and without *L. maculans* infection are difficult to obtain in controlled conditions, by contrast to other pathosystems, such as *Z. tritici* on wheat residues (Kerdraon et al., 2019b) or *Plasmodiophora brassicae* on *B. napus* (Lebreton et al., 2019). There is also, generally, a large difference between the microbial communities present on plants in field conditions and those obtained in greenhouses (Ritpitakphong et al., 2016). The use of fungicides, another plausible technical solution for achieving healthy conditions in the field, would have affected non-targeted endophytic or pathogenic fungal communities (Karlsson et al., 2014; Knorr et al., 2019). While host genetics has been shown to influence the plant and root microbiome strongly (Rybakova et al., 2017; Sapkota et al., 2015; Wagner et al., 2016), the use of near-isogenic lines differing only by the presence or absence of a specific resistance gene has limited the ‘genotype’ effect (Newton et al., 2010). This strategy was effective as it made it possible to achieve different levels of *L. maculans* on the two types of residues, mimicking infected and healthy conditions, and could be adapted to achieve similar objectives with other pathosystems having a complex biological cycle. However, it was not possible, with this strategy, to distinguish between the effects of the presence/absence of *Rlm11* on the residue microbiome, and those of the presence/absence of *L. maculans*, which was itself strongly affected by *Rlm11*. To our knowledge, this study is the first to investigate the effect of a fungal plant pathogen on microbial communities on near-isogenic lines (Newton et al., 2010). In addition, very few studies have focused on the micro-organisms associated with *L. maculans* during the necrotrophic and saprophytic stages of the life cycle of this pathogen (Kerdraon et al., 2019c; Naseri et al., 2008).

The proportion of *L. maculans-*infected samples was smaller for Darmor-*Rlm11* residue samples than for Darmor residue samples. The detection of *L. maculans* in a few plants assumed to be resistant cannot be explained by contamination with plants lacking *Rlm11*, as the purity of the Darmor-*Rlm11* seed lot was established before the experiment. The frequency of *avrLm11* (virulent) isolates in the local population was tested by pathotyping a population of *L. maculans* collected from the residues used to reinforce the inoculum. None of the 92 isolates tested was virulent on Darmor-*Rlm11*, suggesting that the proportion of virulent isolates in the field ranged from 0% to 3.9% (Daudin et al., 2001). This is consistent with previous epidemiological surveys indicating that less than 4% of *L. maculans* isolates are virulent against *Rlm11* in French pathogen populations (Balesdent et al., 2013). These elements suggest that rare virulent isolates may lead to the infection of Darmor-*Rlm11* in field conditions, and account for the lower levels of *L. maculans* colonisation on Darmor-*Rlm11* residues than on Darmor residues, as established by metabarcoding and qPCR.

Fungal community structure clearly differed between Darmor and Darmor-*Rlm11* in July, but not on subsequent sampling dates. This suggests that the presence/absence of the *Rlm11* gene, or the whole introgression, influences the microbiome of the stem base whilst the plant is alive, but does not lead to differential colonisation of the residues by fungi, partly from the bulk soil, later on. Bacterial community structure was slightly affected by the presence of *Rlm11*. In July, the divergences between communities observed including an under-representation of *Pyrenopeziza* (probably *P. brassicae*) in Darmor-*Rlm11*. Darmor is a cultivar known to be susceptible to the fungal pathogen *P. brassicae*, which is prevalent mostly in western France and the north of the UK (Gilles et al., 2000). Several hypotheses can be put forward to account for this unexpected difference between the two near-isogenic lines: (1) the presence of *L. maculans* facilitates *P. brassicae* infection; (2) the resistance of Darmor-*Rlm11* to *L. maculans* isolates carrying the avirulence gene *AvrLm11* gene indirectly prevents the early infection of *B. napus* leaves with *P. brassicae*, with a significant impact on the development of polycyclic light leaf spot epidemics; (3) the introgression of *Rlm11* from *Brassica rapa* to *B. napus* was accompanied by the introgression of a locus conferring resistance to *P. brassicae* or replacing a susceptibility locus initially present in cultivar Darmor (Régine Delourme, INRA IGEPP, comm. pers.). These three hypotheses should be tested in future experiments.

All the fungal genera and most of the most abundant bacterial genera identified here were also detected on oilseed rape residues in a previous study (Kerdraon et al., 2019c). The immediate proximity of the experimental plots of the two studies may explain these similarities. As previously established, bacterial and fungal communities changed with the degradation of residues over the interepidemic period. For example, *Chaetomium*, detected on the residues between July and October, has been shown to colonise oilseed rape residues two months after deposition on the ground (Kerdraon et al., 2019c; Naseri et al., 2008). *Alternaria spp*. accounted for a large percentage of the microbes isolated from infected oilseed rape residues in a previous study (Kerdraon et al., 2019c; Naseri et al., 2008). Similarly, *Alternaria* ASVs had a high RA in this study. The genus *Monographella*, detected on the residues in this study, was previously isolated from the rhizosphere of senescent oilseed rape plants and has been described as an antagonist of *Verticillium* (Berg et al., 2005), but no significant direct interactions between *Verticillium* and *Monographella* were detected by ENA in this study. The existence, or lack of existence, of interaction between species or genus as revealed by ENA can help to identify potential biocontrol agents, but a subsequent validation based on functional testing is required.

The dual approach based on LDA and ENA used here detected an effect of the presence of *L. maculans* on the whole residue microbiome. Based on a similar study focusing on wheat residues hosting *Z. tritici* (Kerdraon et al., 2019b), LDA and ENA were also expected to facilitate the identification of beneficial micro-organisms against residue-borne pathogens, such as *L. maculans*. The stem canker caused by *L. maculans* is usually controlled by the deployment of resistant cultivars, so few studies have focused on interactions between *L. maculans* and potential biocontrol agents. We detected a few direct interactions between *L. maculans* and other micro-organisms in the residue microbial community. These rare, but significant interactions concerned both fungi (*Cryptococcus* in July, *Alternaria* in July and February; *Monographella* in October and February; *Chaetomium* in October), and bacteria (*Nocardioides, Sanguibacter* and *Massilia* in October, *Altererythrobacter* and *Sphingomonas* in December, and *Acidovorax* in February). Interestingly, *Chaetomium* was previously shown to be a biocontrol agent against several pathogenic micro-organisms, including *Sclerotinia sclerotiorum* in oilseed rape (Zhao et al., 2017). These differential interactions may also be related to some of the properties of *L. maculans*, which produces phytotoxins such as sirodesmin PL (Rouxel et al., 1988) that were shown to have *in vitro* inhibitory effects on some fungi and bacteria, while *P. biglobosus* does not produce this toxin (Pedras & Biesenthal, 2000; Elliott et al., 2007). However, fungi usually produce a number of toxic metabolites, and, apart the lack of production of sirodesmin PL by *P. biglobosus*, both species produce a complex set of secondary metabolites whose toxic effects have not been analysed yet (Grandaubert et al., 2014).

This study revealed a key role of *P. biglobosus* in stem residue communities. This fungal species has received much less attention than *L. maculans*, to which it is related. These two pathogenic species belong to the same species complex, but little is known about their effective interaction in plants. The species follow the same infectious cycle, but differ slightly in terms of their ecological niches on oilseed rape, with *L. maculans* colonising the stem base, whereas *P. biglobosus* is restricted to upper parts of the stem (Fitt et al., 2006a). In Europe *P. biglobosus* is considered to be much less important than *L. maculans*. A key finding of the metabarcoding analysis was the higher RA for *Plenodomus* than for *Leptosphaeria*. By contrast, other data acquired in the same experimental area and the same cropping season (2016-2017) (Kerdraon et al., 2019c) highlighted similar RA values for *Leptosphaeria* (*L. maculans*) and *Plenodomus* (*P. biglobosus*) on residues of cv. Alpaga, which is susceptible to *L. maculans*. The lower RA of *Leptosphaeria* on cv. Darmor than on cv. Alpaga can be attributed to the high level of quantitative resistance of cv. Darmor to *L. maculans*. Thus, when Darmor or Darmor-*Rlm11* stems are infected with low levels of *L. maculans, P. biglobosus* freely colonises the stem base. The high RA of *Plenodomus* in cv. Darmor also suggests that the high level of quantitative resistance of cv. Darmor is not effective against *P. biglobosus*. The hypothesis that *P. biglobosus* colonises the stem base niche in the absence of *L. maculans* or in the presence of low-level *L. maculans* colonisation is consistent with the results of ENA, which detected no direct interactions between *Leptosphaeria* and *Plenodomus*. This should be also connected to the results of field studies from Australia whereby *P. biglobosus* has been shown to colonise the stems of *B. juncea* plants, a species resistant to most *L. maculans* isolates (Elliott et al., 2011; Van de Wouw et al., 2008).

Overall, the data presented here clearly suggest that the impact of *P. biglobosus* on the development of stem canker symptoms and yield losses may have been underestimated, at least in some parts of Europe, and should therefore be re-evaluated. The most abundant ASV from *Plenodomus* (93.8% of the ASVs for this genus) had a central position in the July networks, in terms of degree and betweenness centrality, thus highlighting the importance of this species in the living plant. By contrast, the non-central position of *Leptosphaeria* in networks suggests a weak impact on the residue microbial community as a whole, and vice versa, despite as the role of this fungus as the causal agent of the disease. These observations may reflect the high level of quantitative resistance to *L. maculans* in cv. Darmor, clearly illustrated by the low RA of *Leptosphaeria* ASVs even in the absence of the *L. maculans* resistance gene *Rlm11*.

This study provides an original example of research revealing alterations to the crop residue microbiome induced by the creative modulation of pathogen levels with a resistance gene, with high-throughput DNA sequencing techniques. The dual approach based on LDA and ENA revealed that the pathogen studied here (*L. maculans*), although prominent, appeared to play a weak role in ecological interaction networks, whereas another initially neglected pathogen (*P. biglobosus*) was found to be a keystone taxon in the networks at harvest. As previously shown in studies of wheat residues hosting *Z. tritici* (Kerdraon et al., 2019b), this approach can help to identify beneficial micro-organisms against residue-borne pathogens. More broadly, it can be used to decipher the role of complex interactions between multi-species pathosystems and other microbial components of crop residues in the shaping of a plant-protective microbiome.

## Experimental procedures

### Oilseed rape field assay

The two cultivars Darmor (INRA-SERASEM, 1984) and its derived isogenic line Darmor-*Rlm11* were sown in 1.75 m × 8 m microplots on the Grignon experimental domain (Yvelines, France; 48°51′N, 1°58′E). The isogenic line Darmor-*Rlm11* was generated by introgression of the resistant gene *Rlm11* from *Brassica rapa* in *Brassica napus* as described by Balesdent et al. (2013). Before sowing, Darmor-*Rlm11* plants were checked as previously described (Balesdent et al., 2013) to confirm the purity of the seed lot. All 167 plants tested possessed *Rlm11*. Seeds were sown in September 2016 with an INOTEC single-seed speeder, at a density of 60 plants per m^2^. Oilseed rape residues of a susceptible cultivar (cv. Alpaga) from the previous growing season (2015-2016) were placed on the ground one month later (October 17, 2016), with the stubble from 50 plants added per plot, to increase inoculum pressure (Brun et al., 2010). We checked for the presence of typical leaf lesions on plants three times between November 14 and December 2, 2016. The first lesions appeared on the leaves in early December, but disease pressure was too low for a quantitative assessment of the primary inoculum at the rosette stage. One month before full maturity (June), 60 plants of each plant genotype were collected from each plot to establish the ‘G2 score’ (Aubertot et al., 2004) characterising stem canker severity. Briefly, each plant was cut at the collar and disease severity (proportion of tissue necrotic due to *L. maculans*) was estimated and converted into a score from 1 (no necrosis) to 6 (100% of the section necrotic). The G2 score, ranging from 0 (no disease) to 9 (all plants lodged), was then calculated for each plot, as described by Aubertot et al. (2004).

### Preparation of oilseed rape residues

After harvest (July), 60 plants of Darmor and 60 of Darmor-*Rlm11* were collected. The plants were washed and portions of the stem were cut from 3 cm below and 6 cm above the crown to serve as ‘residues’. We first used residues from 15 Darmor and 15 Darmor-*Rlm11* plants to characterise the microbial communities present at harvest. We then weighed the residues from the remaining 45 plants of each cultivar and placed them in nylon bags (residues from one plant par bag). In late July, the bags were placed on the ground in a neighbouring plot in the same experimental area (wheat-oilseed rape rotation (Kerdraon et al., 2019c)) at 15 sampling points 20 m apart, with 3 Darmor and 3 Darmor-*Rlm11* bags per sampling point.

The impact of season on the fungal and bacterial communities of the residues was assessed by collecting the residues from one Darmor and one Darmor-*Rlm11* bag on each of three dates (October, December and February) from each sampling point (15 residue replicates per sampling date and per cultivar). The residues were rinsed with water, dried in air, and weighed to assess degradation. Residues were then crushed individually in liquid nitrogen with a zirconium oxide blender in a Retsch(tm) Mixer Mill MM 400 for 60 seconds at 30 Hz.

### DNA extraction, PCR and Illumina sequencing

Total DNA was extracted with the DNeasy Plant Mini kit (Qiagen, France), according to the manufacturer’s instructions, with minor modification as described by Kerdraon et al. (2019b). The Internal Transcribed Spacer 1 (ITS1) genomic region and the v4 region of the 16S rRNA gene were amplified for the analysis of fungal and bacterial community profiles, respectively. Two rounds of amplifications were performed following the standard operating procedures for building amplicon libraries. The first round of amplifications (30 cycles of PCR) was performed with the ITS1F/ITS2 (Buée et al., 2009) and 515f/806r (Caporaso et al., 2011) primers and the Type-it® Microsatellite PCR kit, as described by Kerdraon et al. (2019b). The amplicons were then sent to an external sequencing platform (Genomics and Transcriptomics GeT core facility of Genotoul, Toulouse, France) for the second round of amplification (12 cycles) performed with primers containing Illumina adapters and indices (Kerdraon et al., 2019b). Libraries were sequenced in one run, with MiSeq reagent kit v3 (600 cycles).

### Quantification of *L. maculans* and *P. biglobosus* by quantitative real-time PCR (qPCR)

We first checked whether, under the conditions of our study, there was a quantitative relationship between biomass and the number of sequencing reads generated by metabarcoding analysis (Lamb et al., 2019). We quantified the DNA of *P. biglobosus* and *L. maculans* by qPCR in 33 of the residue samples used for our metabarcoding analysis (same DNA used for both experiments). For the quantification of *L. maculans* (LM assay), the forward primer LM_EF1-F5 (TGGACACTTCTTCTTGACAA), reverse primer LM_EF1-F5 (TGGACACTTCTTCTTGACAA) and probe LM_P (TACCACGTTCACGCTCGGCC) were designed based on the partial sequence of the EF1 alpha gene. For the quantification of *L. biglobosa* (LB assay), the forward primer LB_ACT_F1 (TTGAGAGCGGTGGCATCCA), reverse primer LB_ACT_R2 (CACCAGACTGTGTCTTTGTC) and probe LB_P (ACGATGTTGCCGTAGAGGTCTTTC) were designed based on the partial sequence of the actin gene. For each assay (LM or LB) the reaction mixture contained 5 µl of sample DNA, 12.5 µl of 2x qPCR MasterMix Plus without UNG (Eurogentec, Belgium), 100 mM probe and 300 mM forward and reverse primers, in a total volume of 25 µl. The quantification cycle (Cq) values were acquired for each sample with the CFX96 Real Time PCR Detection System (Biorad, USA), under the following cycling conditions: 10 min at 95°C for initial denaturation, 40 cycles of amplification consisting of heating at 95°C for 15 s and 1 min at the annealing temperature of 60°C for LM assay and 62°C for LB assay. For the standard calibration curve, reference DNA concentrations from 5 ng to 0.005 ng were analysed in triplicate. A standard curve was generated for each PCR plate. The fluorescence threshold was automatically calculated with Biorad CFX Manager software. The results of the qPCR experiments were compared with the numbers of reads obtained for *Leptosphaeria* or *Plenodomus*. A Spearman’s rank correlation test was performed with the R package ‘car’, to investigate the relationship between the two variables (Fox and Weisberg, 2018).

### Sequence processing

Primer sequences were removed with Cutadapt version 1.8 (Martin, 2011). Fastq files were processed with DADA2 version 1.8.0 (Callahan et al., 2016) as recommended in the DADA2 Pipeline Tutorial (1.8) (Callahan), using the following parameters: truncLen = c(200,150), maxN = 0, maxEE = c(1,1), truncQ = 5. A mock sample composed of DNA from known micro-organisms was included in the sequencing run (Suppl. Figure 6) to establish a detection threshold for spurious haplotypes. At a threshold of ≤ 3 ‰ of the library size, amplicon sequence variants (ASV) were considered spurious and were removed from the sample. Taxonomic affiliations of ASVs were performed with a naive Bayesian classifier (Wang et al., 2007) implemented in DADA2. ASV derived from 16S rRNA gene and ITS1 reads were classified at a minimum bootstrapping threshold of 50 with the RDP trainset 14 (Cole et al., 2009) and the UNITE7.1 database (Abarenkov et al., 2010), respectively. We used the default paramater of the ‘assignTaxonomy’ function with minimum bootstrapping threshold of 50. ASVs classified as chloroplasts (for bacteria) or unclassified at the phylum level were also removed from each sample. After quality filtering, the median read numbers were 27,122 and 26,436 for 16S rRNA gene and ITS1 sequences, respectively (Suppl. Figure 1).

### Microbial community analysis

Normalisation by proportion was performed to standardise total library size for each sample. The effects of season (sampling date) and the presence of *Rlm11* on alpha diversity were assessed with the Shannon index, complemented with the observed richness (number of ASVs) and the Pieoule’s index (taxa evenness). The phylogenetic diversity was not estimated because of the polymorphism in size of ITS1 that hampered sequence alignment. The divergences in microbial community composition, illustrated by MDS (ape package v 5.2.; Paradis and Schliep), were assessed with the Bray-Curtis dissimilarity matrix, calculated with the phyloseq package (v 1.24.2; McMurdie and Holmes, 2013). The effects of season and of the presence of *Rlm11* on microbial community structure were investigated by performing PERMANOVA with the ‘margin’ option (ADONIS2 function, ‘vegan’ package; Oksanen et al., 2017). The ‘PairwiseAdonis’ function was used to characterise the divergence between conditions in post-hoc tests (Martinez Arbizu, 2019).

### Influence of the *Rlm11* gene and of the presence of *L. maculans* on the residue microbiome

The impact of the presence of *Rlm11* on the microbial communities was investigated by performing a linear differential analysis (LDA) for each sampling date with LefSe (Segata et al., 2011) implemented in Galaxy (http://huttenhower.org/galaxy). LefSe uses the Kruskal-Wallis rank-sum test with a normalised relative abundance matrix to detect features with significantly different abundances between assigned taxa, and performs LDA to estimate the effect size of each feature. The alpha risk for the Kruskal-Wallis test was set at 0.05 and the effect-size threshold was set at 2.0.

For the identification of interactions between *L. maculans* and other organisms, ecological interaction analysis (ENA) was performed for each time point, for both cultivars (Darmor and Darmor-*Rlm11* samples considered together). Networks were computed with SPIEC-EASI (Kurtz et al., 2015), from combined bacterial and fungal datasets (Tipton et al., 2018) for each sampling date (30 samples per network). Network sensitivity was increased by removing rare ASVs (Berry and Widder, 2014), by defining a minimum threshold of six occurrences. The Meinshausen and Bühlmann (MB) method was used as graphical inference model for the four networks (Kurtz et al., 2015). The StARS variability threshold was set at 0.05. The betweenness centrality of each node and node degree were estimated with the igraph package (v. 1.2.2.; Csardi and Nepusz).

The two analyses — LDA and ENA — were combined by constructing subnetworks focusing on the interactions between ASVs identified as differential in Darmor relative to Darmor-*Rlm11* by the LDA approach. The differential ASVs highlighted by LDA and their adjacent nodes were used for the analysis of subnetworks constructed with Cytoscape (v. 3.6.1; Shannon et al., 2003). The scripts for network construction and analysis are available from GitHub.

## Acknowledgements

This study was supported by a grant from the *European Union Horizon Framework 2020 Program* (*“EMPHASIS Project”*, Grant Agreement no. 634179) covering the 2015- 2019 period. It was performed in collaboration with the GeT core facility, Toulouse, France (http://get.genotoul.fr/), supported by *France Génomique National Infrastructure*, funded as part of the “*Investissement d’avenir*” program managed by *Agence Nationale pour la Recherche* (contract ANR-10-INBS-09). We thank Régine Delourme and Pascal Glory (INRA, UMR IGEPP) for providing seeds of the near-isogenic *B. napus* lines, Angelique Gautier (INRA, UMR BIOGER) for preparing the mock samples, Noémie Jacques for performing qPCR experiments, Christophe Montagnier (INRA Experimental Unit, Thiverval-Grignon) for managing the field trial, and Julie Sappa for her help correcting our English.

## Data availability statement

The raw sequencing data are available from the European Nucleotide Archive under study accession number PRJEB35369 (samples ERS4024677 to ERS4024484). We provide the command-line script for data analysis and all necessary input files via GitHub (https://github.com/LydieKerdraon/2019_OilseedRape).

## Authors’ contributions

LK, VL, FS, MHB, MB conceived the study, participated in its design, and wrote the manuscript. LK conducted the experiments and analysed the data. FS and VL supervised the project. All the authors read and approved the final manuscript.

## Supporting information

**Supplementary Figure 1.**
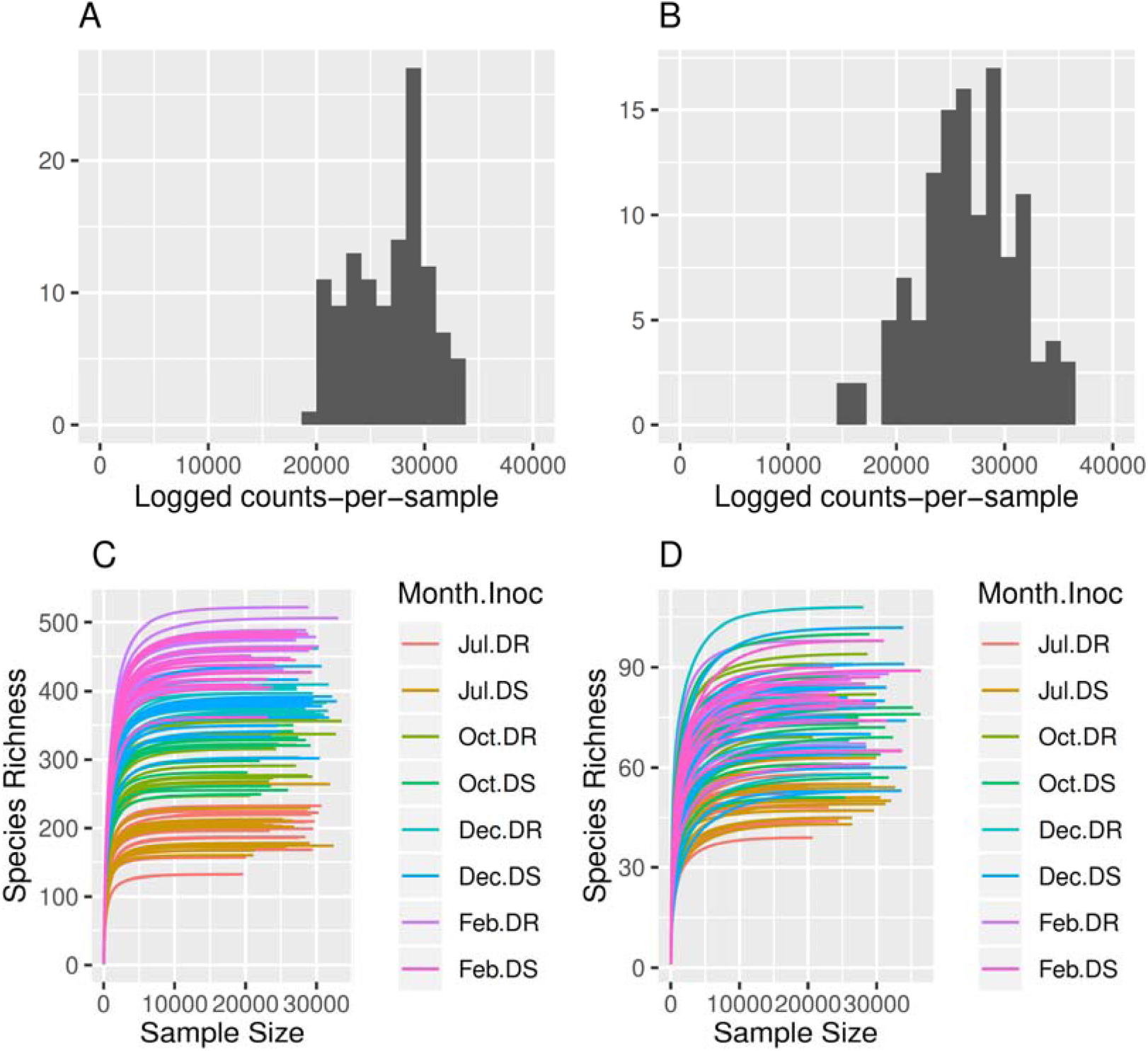
Number of bacterial reads for 16S rRNA gene (A) and of fungal reads for ITS1 (B) after quality filtering. Rarefaction curves obtained for bacteria (C) with 16S rRNA gene and for fungi (D) with ITS1 sequences.

**Supplementary Figure 2.**
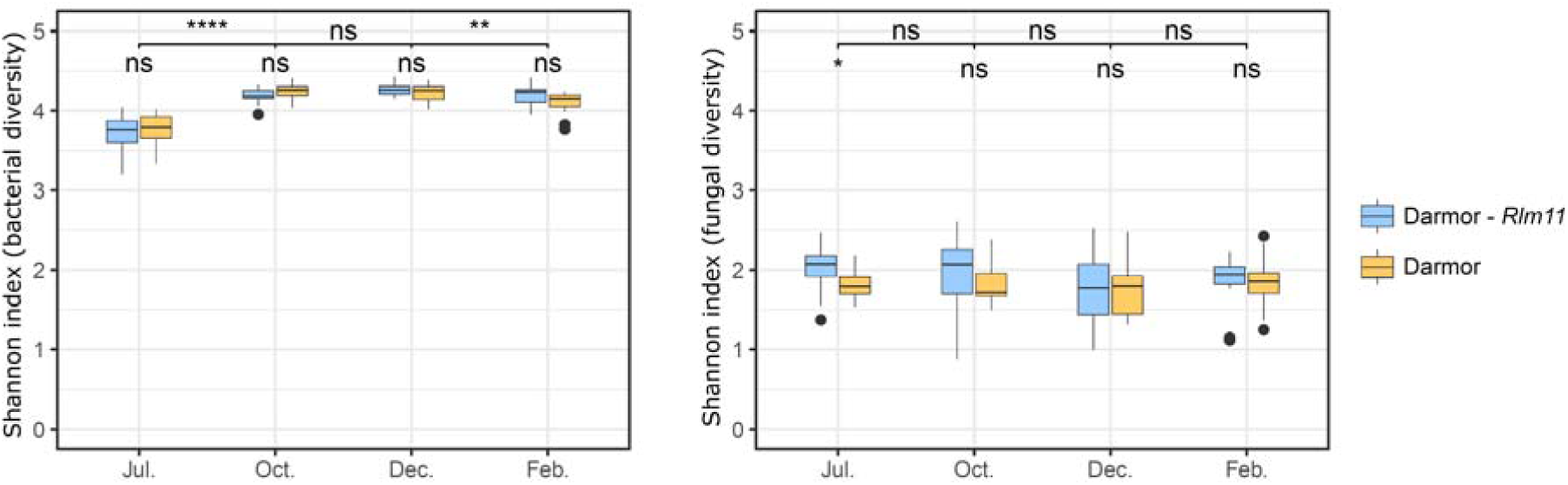
Alpha diversity of microbial assemblages associated with oilseed rape residues. Each box represents the distribution of Shannon index for 15 sampling points per treatment. Wilcoxon tests (NS: not significant; * *p*-value < 0.05; ** *p*-value < 0.01; *** *p*- value < 0.001) were performed for host cultivar (Darmor, Darmor-*Rlm11*) and sampling date (July, October, December, February).

**Supplementary Figure 3.**
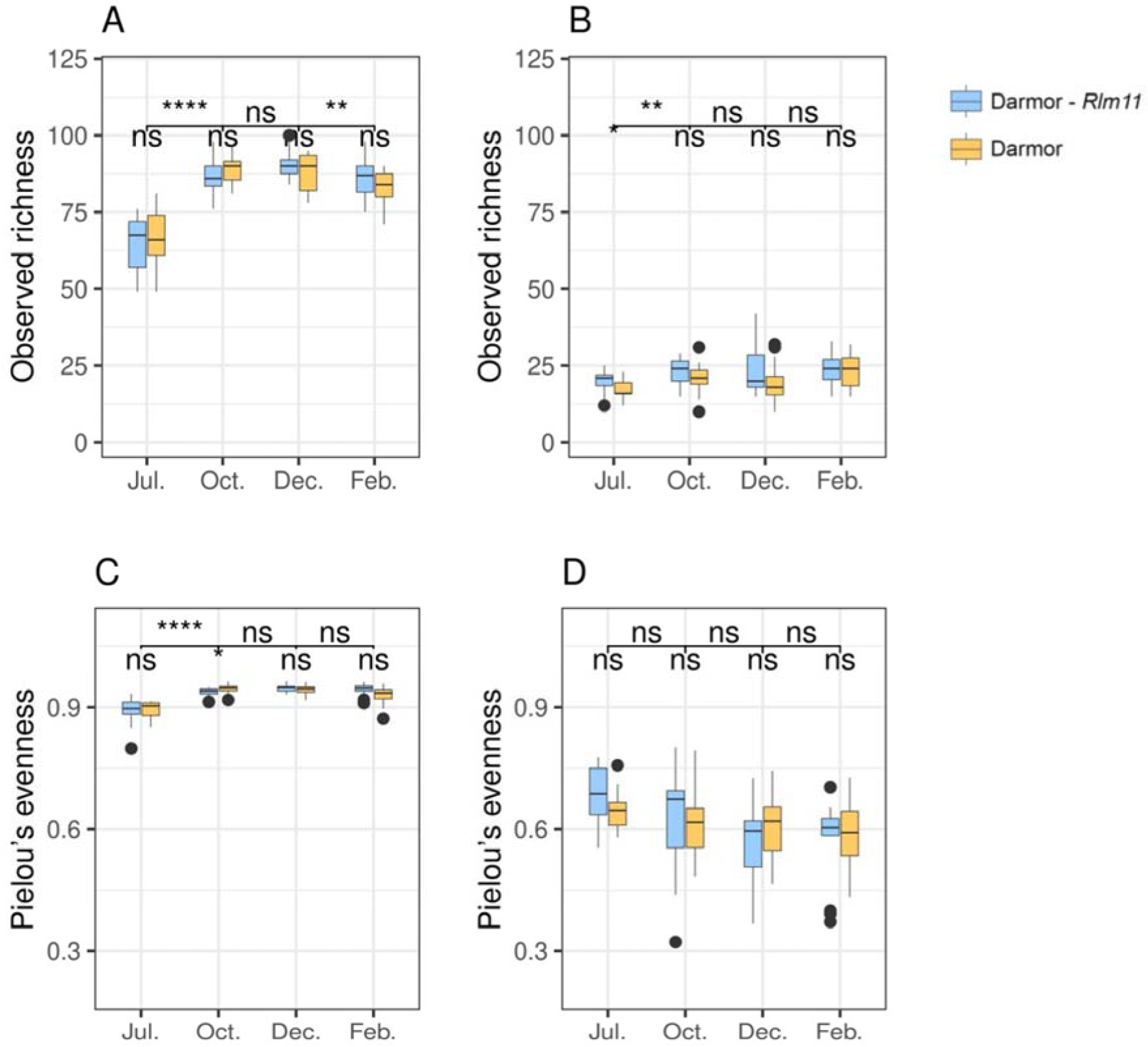
Observed richness and taxa evenness of microbial assemblages associated with oilseed rape residues. Observed richness (number of ASVs) was estimated for bacteria (A) with 16S rRNA gene and for fungi (B) with ITS1 sequences. Taxa evenness (Pielou’s index) was estimated similarly for bacteria (C) with 16S rRNA gene and for fungi (D) with ITS1 sequences.

**Supplementary Figure 4.**
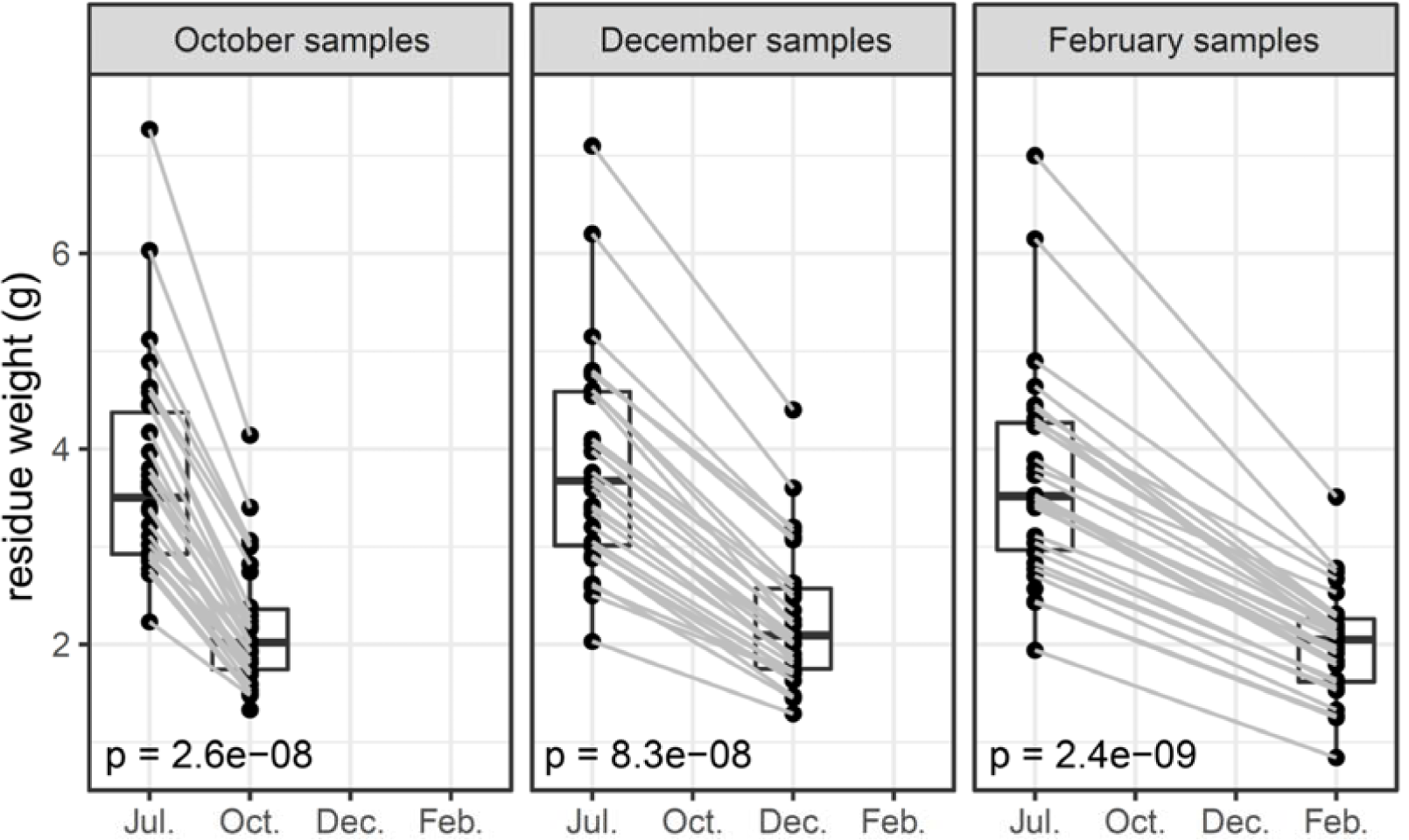
Degradation of oilseed rape residues over time. Each set of residues was weighed in July before being placed in nylon bags, and then again at each sampling date. The grey lines indicate the decrease in weight of the set of residues from July to each sampling date. Wilcoxon tests were performed to assess the significance of the weight decrease and the *p* value is indicated on each graph.

**Supplementary Figure 5.**
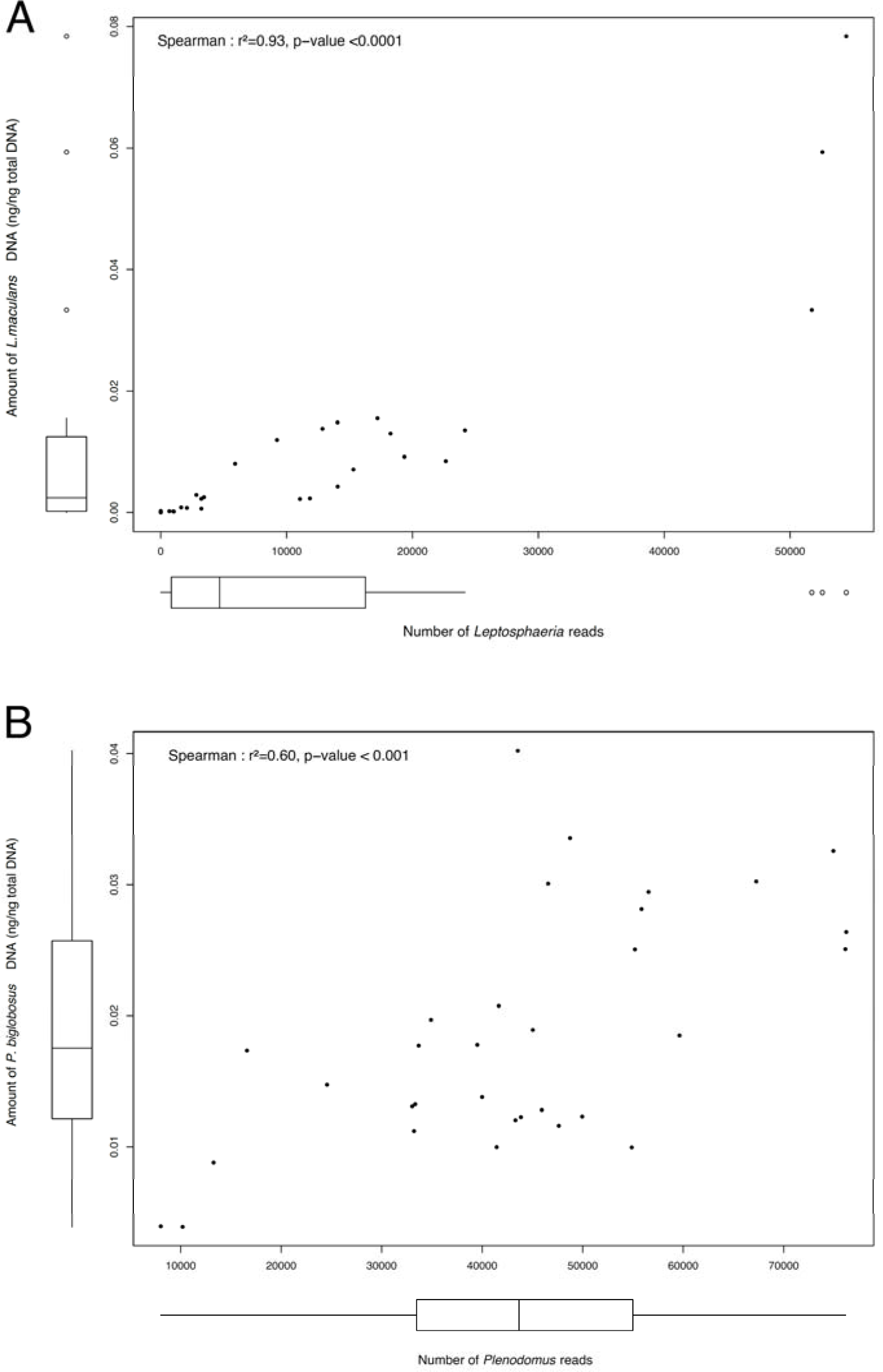
Relationship between the amount of (A) *Plenodomus biglobosus* and (B) *Leptosphaeria maculans* DNA quantified by qPCR (in ng/ng of total DNA) and the sequence read numbers obtained by metabarcoding with a set of 33 oilseed rape residue samples. A Spearman’s rank correlation test (ρ; p-value) was performed with the R package “car” to evaluate this relationship. DNA quantification with qPCR was performed on ‘*Leptosphaeria biglobosa*’ whereas metabarcoding results led to the identification of ‘*Plenodomus*’ but not of the corresponding species. For the sake of consistency, a single name, ‘*P. biglobosus*’, is used in B.

**Supplementary Figure 6.**
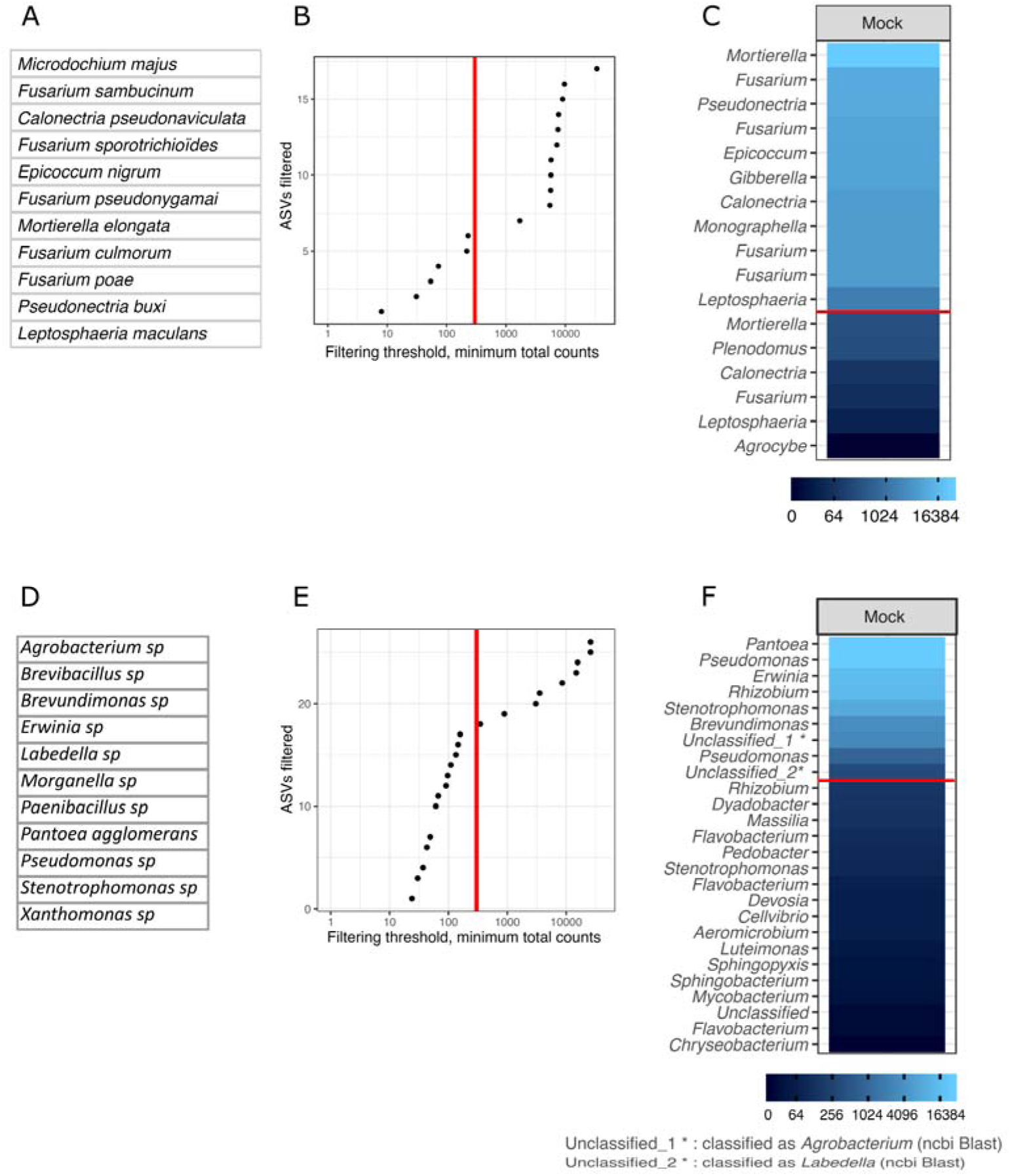
Mock sample analysis for filtering on the relative abundance of ASVs of the fungal (A, B, C) and bacterial (D, E, F) sequencing runs. (A, D) Composition of mock samples. All microbial DNAs were pooled at equimolar concentrations. (B, E) Filtering on the relative abundance of ASVs. Library size was normalised by proportion before analysis. All samples were sequenced in the same run, and a threshold (red line) was set at 3 ‰ for both bacterial and fungal community analyses. (C, F) ASVs detected in each mock sample. The name of the ASV corresponds to the taxonomic genus to which it is affiliated. All genera present in fungal mock samples were detected (*Microdochium majus* and *Fusarium nivale* are synonymous, *Monographella* is the teleomorph of *Microdochium*/*Fusarium, Gibberella* and *Fusarium* are synonymous), but some bacterial genera were not detected in bacterial mock samples. The red line corresponds to the threshold of 3 ‰ of the size of the library.

